# N-terminal signals in the SNX-BAR paralogs Vps5 and Vin1 guide coat complex formation

**DOI:** 10.1101/2024.01.24.576755

**Authors:** Shawn P. Shortill, Mia S. Frier, Michael Davey, Elizabeth Conibear

## Abstract

Endosomal coat complexes assemble by incorporating membrane-binding subunits such as those of the sorting nexin (SNX) family. The *S. cerevisiae* SNX-BAR paralogs Vin1 and Vps5 are respective subunits of the endosomal VINE and retromer complexes that arose from a fungal whole genome duplication. Interactions mediated by the Vin1 and Vps5 BAR domains are required for protein complex assembly and membrane association. However, a degree of promiscuity is predicted for yeast BAR-BAR pairings, suggesting that another mechanism guides the formation of specific endosomal coat complexes. Previous work by our group and others has implicated the unstructured N-terminal domains of Vin1 and Vps5 in complex assembly. Here, we map N-terminal signals in both SNX-BAR paralogs that contribute to the formation and function of two distinct endosomal coats *in vivo*. Whereas Vin1 leverages a polybasic region and adjacent hydrophobic motif to bind Vrl1 and form VINE, the N-terminus of Vps5 interacts with the retromer subunit Vps29 at two separate sites. We show that one of these Vps5 motifs binds to a conserved hydrophobic pocket in Vps29 that is shared with other accessory proteins and targeted by a bacterial virulence factor in humans. Lastly, we examined the sole isoform of Vps5 from the milk yeast *K. lactis* and found that ancestral yeasts may have used a nested N-terminal signal to form both VINE and retromer. Our results suggest that the specific assembly of Vps5-family SNX-BAR coats depends on inputs from unique N-terminal sequence features in addition to BAR domain coupling, expanding our understanding of endosomal coat assembly mechanisms.

## Introduction

SNX-BARs are a conserved subfamily of dimerizing sorting nexin proteins that deform membranes into cargo-enriched tubules to promote membrane transport (Frost et al., 2009; Shortill, Frier, & Conibear, 2022; van Weering et al., 2010). Many SNX-BARs contain an extended, unstructured N-terminus, a lipid-binding PX domain and a concave BAR domain that drives dimerization (Ma & Burd, 2020; Van Weering & Cullen, 2014). Partner selection is thought to be enforced by complementary sets of charged amino acids in the extended interfaces of BAR domains, establishing a pairing code that promotes the specific assembly and function of SNX-BARs (Van Weering et al., 2012). Mutation of either the internal matching charged residues or the distal ends of BAR domains that facilitate tip-to-tip SNX-BAR oligomerization and tubule extension disrupts sorting (Dislich et al., 2011; Lopez-Robles et al., 2023; Van Weering et al., 2012).

The *S. cerevisiae* SNX-BAR paralogs Vps5 and Vin1 arose from a fungal whole genome duplication event (Byrne & Wolfe, 2005; Wolfe & Shields, 1997) and subsequently diverged to assume new roles. We previously determined that Vin1 specifically binds to the VPS9-domain GEF Vrl1 to form a novel endosomal coat complex that we named VINE (Shortill et al., 2022). In contrast, Vps5 is a well-studied subunit of the ubiquitous retromer complex—a conserved endosomal assembly that promotes the retrograde transport of numerous protein cargoes (Bean et al., 2017; Burd & Cullen, 2014; Ma & Burd, 2020). Yeast retromer has been described as a heteropentamer composed of a Vps5-Vps17 SNX-BAR dimer and a Vps26-Vps35-Vps29 trimer (Seaman et al., 1998). In humans retromer refers strictly to the VPS26-VPS35-VPS29 trimer, which can associate with SNX-BAR dimers or other membrane adaptors to regulate diverse sorting processes (Arighi et al., 2004; Wassmer et al., 2007; Cui et al., 2019; Seaman, 2021; Steinberg et al., 2013; Courtellemont et al., 2022; Kvainickas et al., 2017; Simonetti et al., 2019).

Vin1 associates with Vrl1 through its unstructured N-terminus (Shortill et al., 2022). Likewise, both Vps5 and its human ortholog SNX1 interact with the Vps35-Vps29-Vps26 trimer through their low complexity N-termini, although the motifs involved have yet to be described (Seaman & Williams, 2002; Gullapalli et al., 2004). Because Vin1 and Vps5 evolved from a common ancestor, they could have acquired divergent N-termini that dictate protein complex selectivity, and therefore functional specificity.

Using a combination of structural predictions, live-cell imaging, and biochemical assays, we mapped the motifs in the Vps5 and Vin1 N-termini that facilitate binding to Vps29 and Vrl1, respectively. We identified a shared Leu-Phe motif as well as sequence features unique to either paralog. In the closely related yeast *K. lactis*, the sole Vps5 isoform has nested N-terminal motifs for binding to Vrl1 and Vps29, indicating a possible ancient regulatory relationship between VINE and retromer. Taken together, our findings help explain the functional divergence between Vin1 and Vps5 and demonstrate the contributions of unstructured regions in SNX-BAR proteins to the assembly and function of membrane sorting complexes.

## Results

### Vrl1 recognizes polybasic and Leu-Phe signals in the Vin1 N-terminus

Vin1 and its paralog Vps5 leverage N-terminal interactions to form VINE and retromer, respectively. Previously, we found that Vrl1 interacts with the N-terminus of Vin1, but not Vps5, and that this interaction depends on a 20 amino acid (aa) motif in the Vin1 N-terminus (aa 76-95) (Shortill et al., 2022). To identify the N-terminal sequence elements that allow Vrl1 to distinguish Vin1 from Vps5, we examined the predicted interface between Vin1^76-95^ and Vrl1 that was obtained using AlphaFold2 through the ColabFold platform (Jumper et al., 2021; Mirdita et al., 2022); Figure 1A-C). The Vin1 peptide and Vrl1 make extensive contact, with numerous predicted sidechain interactions that span the full Vin1^76-95^ region.

**Figure 1.**
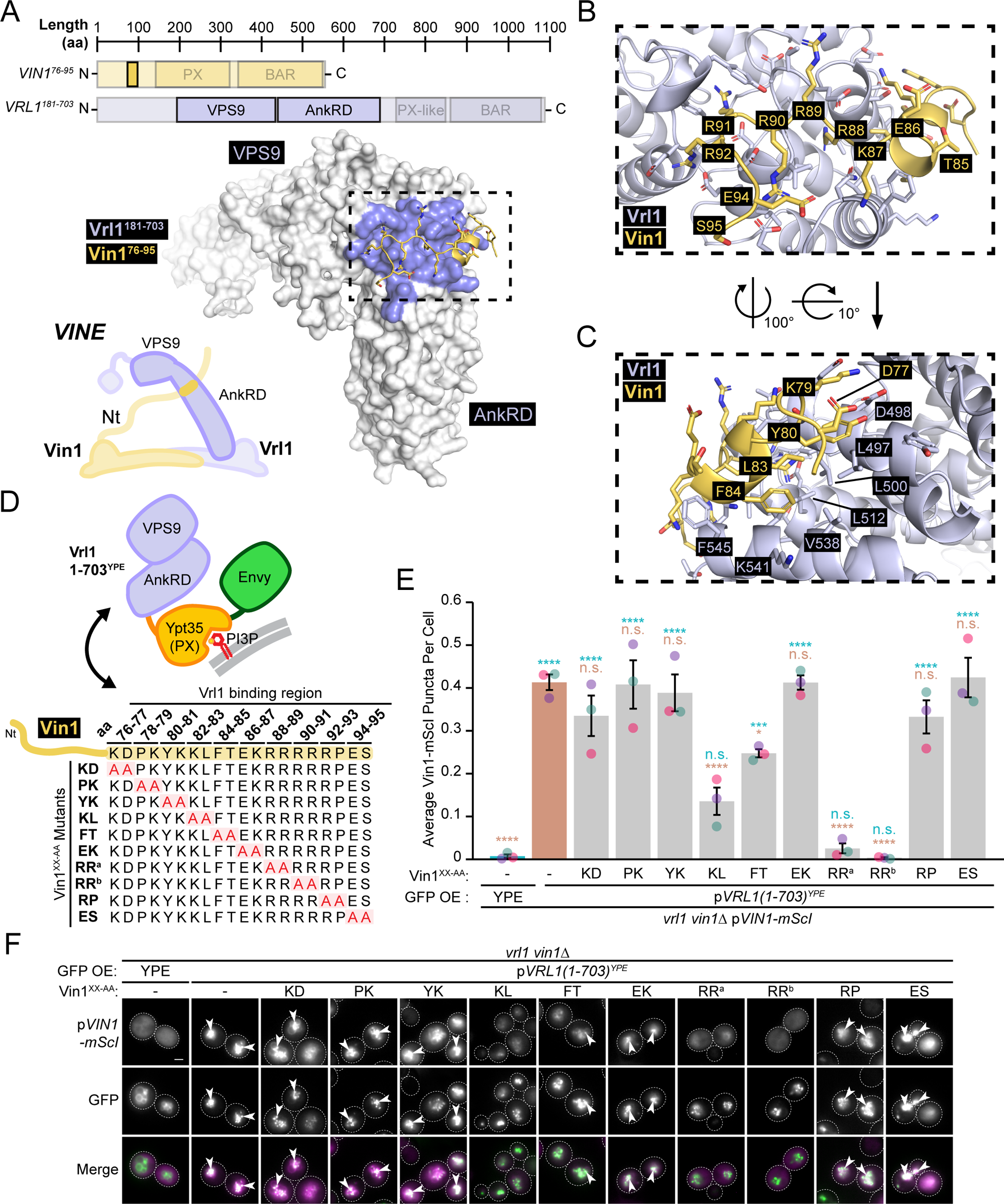
Mapping features in the Vin1 N-terminus that are required for recognition by Vrl1. (A) AlphaFold2-predicted interaction between Vrl1^181-703^ and the Vrl1-binding portion of the Vin1 N-terminus (Vin1^76-95^). Interacting surface on Vrl1 shown in blue. (B) Basic residues within Vin1^76-95^ are predicted to associate with Vrl1. (C) nonpolar residues in Vin1^76-95^ are predicted to associate at an adjacent site on the Vrl1 AnkRD. (D) Diagram of assay used to test recruitment by chimeric Vrl1 construct (top) of the indicated pairwise alanine substitution mutants of the Vin1 N-terminus (bottom). (E) Quantification of RFP puncta per cell for Vin1 mutants. One-way ANOVA with Dunnett’s multiple comparison tests; *n* = 3, cells/strain/replicate ≥ 878; not significant, n.s. = p > 0.05, * = p < 0.05, *** = p < 0.001, **** = p < 0.0001. Blue statistical significance labels correspond to a Dunnett-corrected ANOVA performed against YPE-containing bait while brown labels correspond to a Dunnett-corrected ANOVA performed against wild-type Vin1. (F) Differential effects of pairwise alanine substitution in the Vin1 N-terminus on recruitment by the AnkRD-containing Vrl1(1-703)^YPE^ chimera. Representative images quantified in *E*. Scale bars, 2 µm. Error bars report standard error of the mean (SEM). aa, amino acids. OE, over-expressed. Nt, N-terminus. mScI, mScarletI. RR^a^, R88A R89A. RR^b^, R90A R91A. YPE, Ypt35(PX)-Envy.

We systematically mutated Vin1-mScarletI (mScI) between aa 76-95 by substituting residues with alanine in sets of two (Figure 1D) and determined if these mutants recognize Vrl1 in our previously established chimeric subcellular recruitment assay (Shortill et al., 2022); Figure 1E, F). Mutation of consecutive residues within a Vin1 basic region, R88, R89, R90 and R91 (herein referred to as the “polybasic” site), completely blocked recruitment by the Vrl1 chimera (p<0.0001) while mutation of K82 and L83 or F84 and T85 had more intermediate disruptive effects (p<0.0001 and p<0.05, respectively). This Vin1 polybasic region is predicted to interact with Vrl1 at the conserved acidic patch that we previously demonstrated was necessary for recognition (Shortill et al., 2022); Figure 1B), suggesting that charged interactions at this site serve an important role in VINE assembly.

To further resolve the contribution of the Vin1 K82, L83, F84 and T85 residues to the interaction with Vrl1, we made an additional L83A F84A mutant (Figure 2A) and compared its recruitment by the Vrl1 chimera to that of the previously tested K82A L83A and F84A T85A mutants (Figure 2B, C). Mutation of this Leu-Phe motif caused a severe loss of recruitment by the Vrl1(1-703)^YPE^ chimera (p<0.0001; Figure 2C), suggesting that this pair of hydrophobic residues is critical for VINE formation. The Leu-Phe motif is predicted to interface with a hydrophobic patch in the Vrl1 AnkRD that is adjacent to the acidic site (Figure 1C). Introducing a charged residue to this hydrophobic patch (Vrl1^L497D^) strongly reduced recruitment of the Vin1 N-terminus (Vin1^1-116^-mScI; p<0.0001; Figure 2D, E), suggesting that this is another site of interaction between Vrl1 and the Vin1 N-terminus.

**Figure 2.**
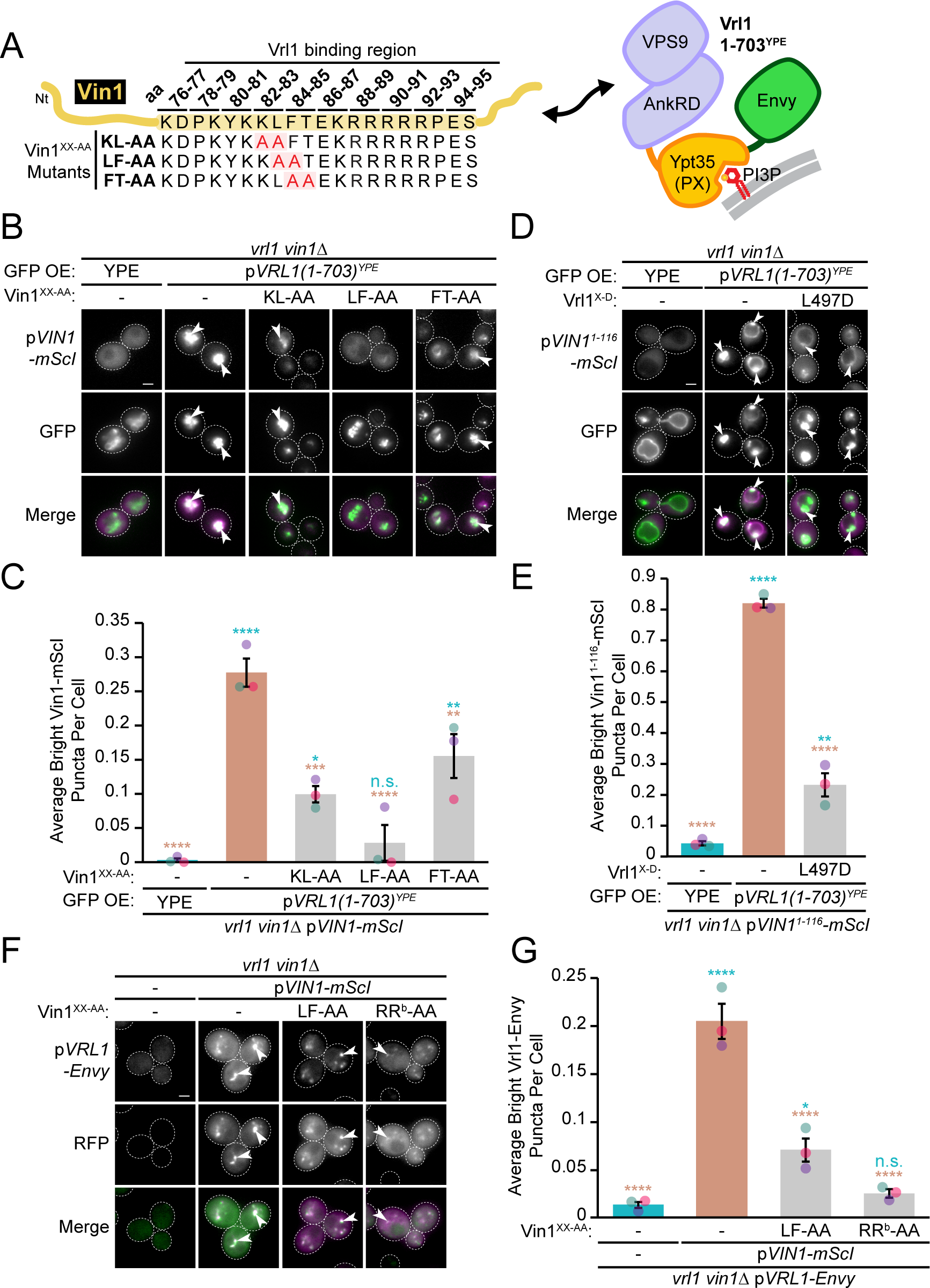
An N-terminal Vin1 Leu-Phe motif is important for recognition by Vrl1 and formation of VINE. (A) Schematic showing the pairwise alanine substitution mutants of Vin1 KL, LF and FT N-terminal residue pairs tested in the chimeric Vrl1 recruitment assay. (B) The Vin1 N-terminal LF-AA mutant displays a severe defect in recruitment by the AnkRD-containing Vrl1(1-703)^YPE^ chimera. (C) Quantification of RFP puncta per cell in *B*. One-way ANOVA with Dunnett’s multiple comparison tests; *n* = 3, cells/strain/replicate ≥ 676; not significant, n.s. = p > 0.05, * = p < 0.05, ** = p < 0.01, *** = p < 0.001, **** = p < 0.0001. Blue statistical significance labels correspond to a Dunnett-corrected ANOVA performed against YPE-containing bait while brown labels correspond to a Dunnett-corrected ANOVA performed against wild-type Vin1. (D) Addition of a charged residue to the Vrl1 hydrophobic patch predicted to interface with the Vin1 N-terminal Leu-Phe motif disrupts recruitment of the Vin1 N-terminus by the AnkRD-containing Vrl1(1-703)^YPE^ chimera. (E) Quantification of RFP puncta per cell in *D*. One-way ANOVA with Dunnett’s multiple comparison tests; *n* = 3, cells/strain/replicate ≥ 863; ** = p < 0.01, **** = p < 0.0001. Blue statistical significance labels correspond to a Dunnett-corrected ANOVA performed against YPE-containing bait while brown labels correspond to a Dunnett-corrected ANOVA performed against wild-type Vrl1. (F) Pairwise substitution of Vin1-mScI L83A F84A or R90A R91A inhibits formation of Vrl1-Envy puncta in *vin1*Δ cells. (G) Quantification of GFP puncta per cell in *F*. One-way ANOVA with Dunnett’s multiple comparison tests; *n* = 3, cells/strain/replicate ≥ 1,108; not significant, n.s. = p > 0.05, * = p < 0.05, **** = p < 0.0001. Blue statistical significance labels correspond to a Dunnett-corrected ANOVA performed against the empty vector *vin1*Δ strain while brown labels correspond to a Dunnett-corrected ANOVA performed against wild-type Vin1-mScI. Scale bars, 2 µm. Error bars report SEM. aa, amino acid. OE, over-expressed. Nt, N-terminus. mScI, mScarletI. RR^b^, R90A R91A. YPE, Ypt35(PX)-Envy.

We also assessed the ability of Vin1 polybasic and Leu-Phe mutants to recruit full-length Vrl1-Envy to puncta. Vrl1 localization in *vin1*Δ cells was restored by Vin1-mScI, and introducing either R90A R91A or L83A F84A mutations to Vin1 disrupted this localization (p<0.0001; Figure 2F, G), suggesting that VINE is only partially assembled and localized in these cells. Taken together, these findings indicate that Vrl1 recognizes two distinct features of the Vin1 N-terminus to form VINE—a polybasic stretch that interacts with an acidic site and a Leu-Phe motif that binds an adjacent hydrophobic patch.

### Vps5 binds to Vps29 through a bipartite motif in its N-terminus

In similar fashion to the interactions that promote formation of VINE, the N-terminus of Vps5 is critical for interaction with the Vps26-Vps35-Vps29 retromer trimer in yeast (Seaman & Williams, 2002). Mutation of a conserved hydrophobic pocket in Vps29 blocks association of the trimer with the Vps5-Vps17 dimer and results in a moderate defect in progression of the vacuolar hydrolase CPY through the retromer-dependent vacuolar protein sorting pathway (Collins et al., 2005). Despite this, how the Vps5 N-terminus associates with the trimer remains incompletely understood.

Using AlphaFold2 (Mirdita et al., 2022), we obtained a confident binding prediction between aa 155-198 of the unstructured Vps5 N-terminus and two distinct sites in Vps29 (Figure 3A, Figure S1A, B). A Vps5 FTDPL motif (herein referred to as “pocket-binding”) was predicted to interact with Vps29 at the previously identified conserved hydrophobic pocket, with Vps5 L196 projecting into the hydrophobic core (Figure 3B). We also identified a Vps5 β-strand composed of the sequence PRILFDS (herein referred to as “sheet-binding”), which is predicted to form a lateral extension of a Vps29 β-sheet found on the surface opposite to the conserved hydrophobic pocket and is separated from the pocket-binding motif by a ∼30 aa unstructured loop region (Figure 3C). Like Vin1, the Vps5 sheet site possesses a Leu-Phe motif predicted to leverage multiple hydrophobic contacts with Vps29 (Figure 1C and 3C).

**Figure 3.**
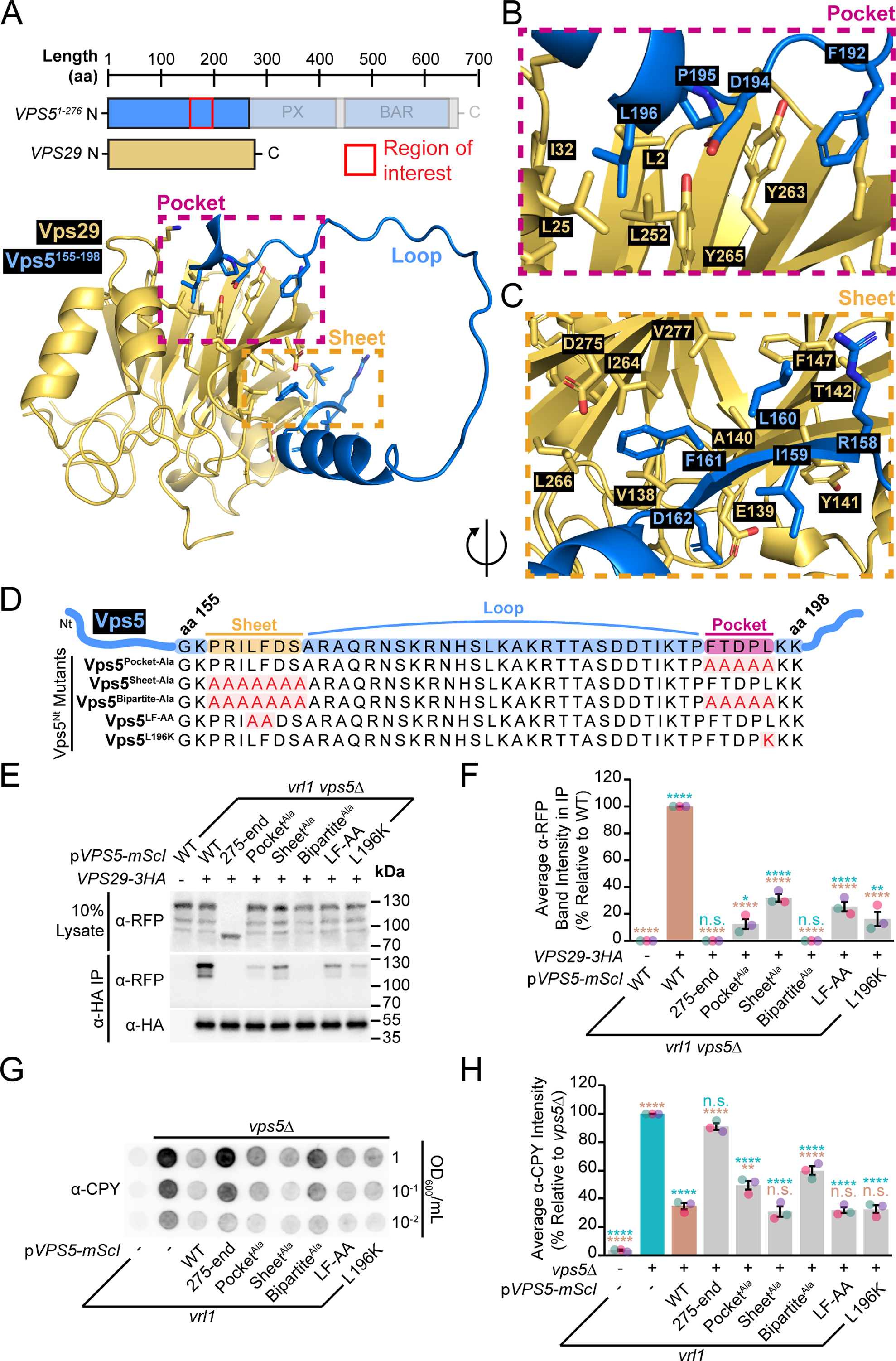
Vps29 binds a bipartite signal in the Vps5 N-terminus. (A) AlphaFold2-predicted interaction of the Vps5 N-terminus (Vps5^1-276^) with Vps29 highlights a bipartite motif between Vps5 aa 155-198. (B) A Vps5 N-terminal FTDPL motif is predicted to interact with Vps29 at a conserved hydrophobic pocket. (C) A Vps5 N-terminal PRILFDS motif is predicted to form a β-strand and extend a Vps29 β-sheet. (D) Diagram of Vps5 substitution mutations in the predicted N-terminal pocket- and sheet-binding sites generated to test retromer assembly and function. (E) Both complete loss and substitution mutagenesis of the Vps5-mScI N-terminus disrupts CoIP with Vps29-3HA. Disruption of either the pocket- or sheet-binding site separately causes a strong binding defect, while combined mutation of both (bipartite-ala) completely blocks binding. (F) Quantification of co-purified Vps5-mScI levels in *E* by densitometry. One-way ANOVA with Dunnett’s multiple comparison tests; *n* = 3; not significant, n.s. = p > 0.05, * = p < 0.05, ** = p < 0.01, **** = p < 0.0001. Blue statistical significance labels correspond to a Dunnett-corrected ANOVA performed against untagged Vps29 while brown labels correspond to a Dunnett-corrected ANOVA performed against 3HA-tagged Vps29 with wild-type Vps5-mScI. (G) Disruption of the Vps5 N-terminal pocket- or sheet-binding sites cause CPY secretion. Paired disruption of the pocket- and sheet-binding sites (bipartite-ala) caused the most severe secretion phenotype. (H) Quantification of secreted CPY by densitometry and normalized to measured secreted CPY from the shown *vps5*Δ strain. One-way ANOVA with Dunnett’s multiple comparison tests; n=3; not significant, n.s. = p > 0.05, ** = p < 0.01, **** = p < 0.0001. Blue statistical significance l abels correspond to a Dunnett-corrected ANOVA performed against a *vps5*Δ strain while brown labels correspond to a Dunnett-corrected ANOVA performed against a *vps5*Δ strain with plasmid-expressed wild-type Vps5-mScI. Error bars report SEM. aa, amino acid. mScI, mScarletI. Nt, N-terminus.

Substituting the pocket- and sheet-binding motifs with alanine in Vps5-mScI either separately or together (bipartite-Ala; Figure 3D) reduced its co-purification with triple hemagglutinin (3HA)-tagged Vps29 (p<0.0001; Figure 3E, F). Disruption of either pocket- or sheet-binding motifs caused intermediate binding defects, whereas combined disruption of both sites resulted in binding loss similar to a full N-terminal truncation mutant (Vps5^275-end^; p<0.0001; Figure 3E, F). Moreover, mutating either the sheet-binding L160 and F161 residues to alanine, or substituting the pocket-binding L196 to lysine, resulted in Vps29 binding defects comparable to complete alanine substitution of either motif (Figure 3D, E, F), suggesting that these residues make important contributions to the interaction with Vps29.

To test the impact of these mutations on retromer-mediated sorting we used a functional assay based on the sorting of newly synthesized CPY at the Golgi. The CPY receptor Vps10 is recognized by retromer at endosomes and recycled to perform additional rounds of CPY transport (Seaman et al., 1998). If recycling is disrupted, soluble CPY is secreted from the cell. Truncation of the Vps5 N-terminus causes CPY processing defects (Seaman & Williams, 2002), and we found that a full N-terminal deletion mutant (Vps5^275-end^) also displays a strong secretion phenotype (Figure 3G, H). The Vps5 N-terminal pocket- and sheet-binding mutants secreted CPY at levels roughly proportional to the severity of demonstrated Vps29 binding deficiency, although not to the same level as truncation of the N-terminus (p<0.0001; Figure 3E, F). Taken together, these data suggest that Vps5 interacts with Vps29 through an N-terminal bipartite pocket- and sheet-binding motif that is critical for both retromer assembly and efficient sorting.

Because both Vps5 and Vin1 leverage their N-termini to form retromer and VINE, respectively, we wondered if these sites could act as sole determinants of coat formation. To test this, we generated a series of mScI-tagged truncation mutants of Vin1 and Vps5 containing either the N-terminus or PX-BAR regions alone, as well as chimeras with respective N-termini swapped to that of the paralog (Figure S2A). None of the Vin1-mScI fragments or Vin1^Nt^ chimera were able to recover the punctate localization of Vrl1-Envy in *vin1*Δ cells (Figure S2B), suggesting that Vrl1 requires input from both the Vin1 N-terminus and Vin1 BAR domain to form VINE. Conversely, while both the Vps5 N-terminus alone and Vps5^Nt^-Vin1^ΔNt^ chimera failed to recover the punctate localization of the retromer subunit Vps35-GFP in *vps5*Δ cells, the Vps5 PX-BAR region alone was able to partially complement (Figure S2C), in line with a previous report (Seaman & Williams, 2002). Taken together, these findings indicate that, while the N-termini of these SNX-BAR paralogs are important for determining coat complex membership, they are not sufficient and input from the respective SNX-BAR domains also contributes to selectivity.

### The sole Vps5 protein from milk yeast *K. lactis* forms both VINE and retromer

In *S. cerevisiae*, protein products of the duplicated *VIN1* and *VPS5* genes have diverged to assume separate roles in forming the VINE and retromer complexes, respectively. Ancestors of the modern milk yeast *K. lactis* did not undergo the same whole genome duplication (WGD) as those of *S. cerevisiae* (Kellis et al., 2004; Keogh et al., 1998), and therefore its genome encodes a single ortholog of the *VPS5* gene. Interestingly, the *K. lactis* genome also encodes orthologs of both the *VRL1* and *VPS17* genes (Byrne & Wolfe, 2005), suggesting that the single copy of Vps5 in this organism may form both VINE and retromer (Figure 4A).

**Figure 4.**
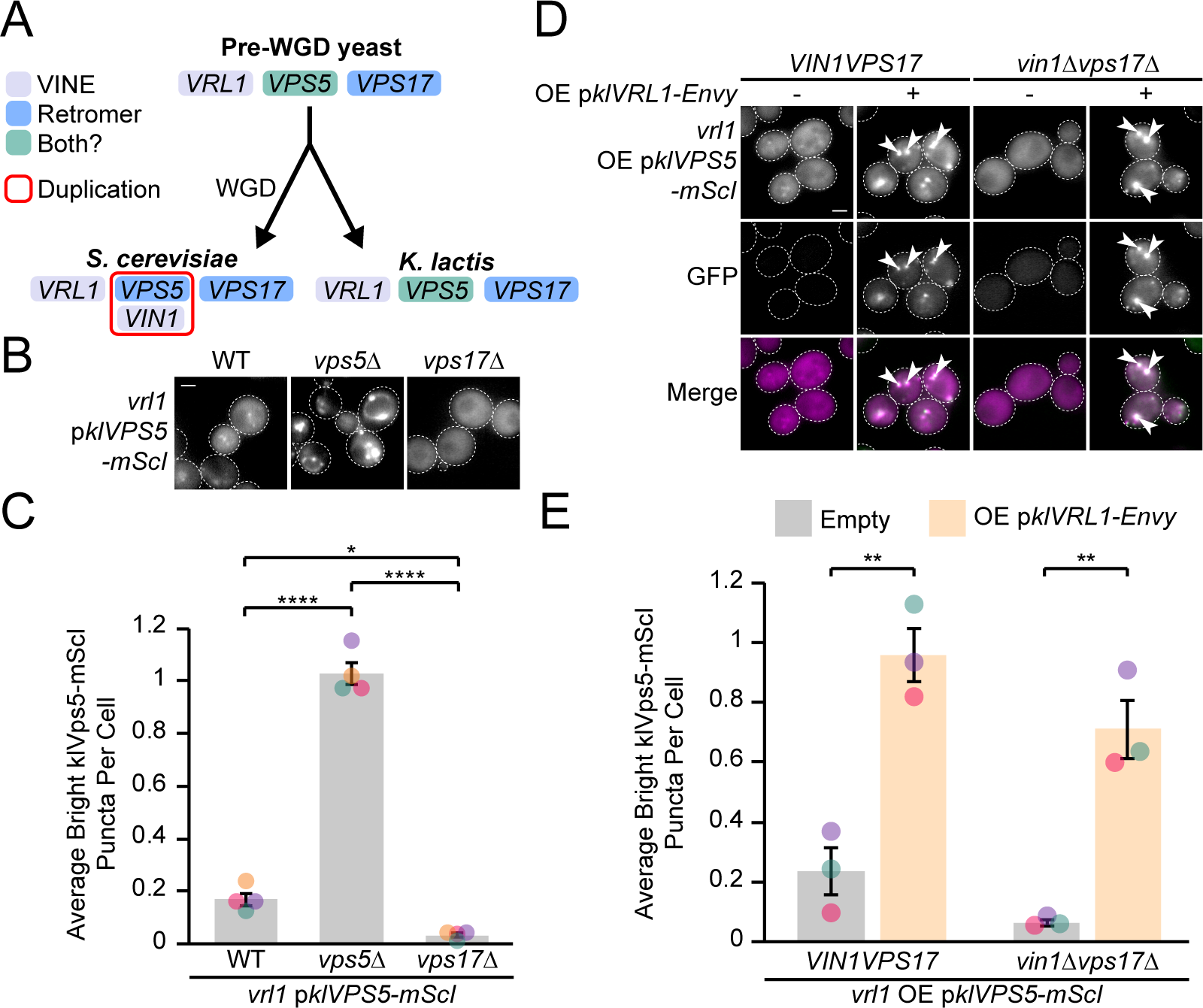
The single *K. lactis* Vps5 isoform forms both retromer and VINE. (A) Diagram of Vps5 duplication status in *S. cerevisiae* and *K. lactis* resulting from an ancient whole genomic duplication event. The single copy of Vps5 in *K. lactis* may bind both Vrl1 and Vps17 to form VINE and retromer, respectively. (B) Ectopically expressed *K. lactis* Vps5 (klVps5-mScI) localizes to puncta in *S. cerevisiae*. Localization depends on Vps17 and is enhanced in a *vps5*Δ strain. (C) Quantification of RFP puncta per cell in *B*. One-way ANOVA with Tukey’s multiple comparison tests; *n* = 3, cells/strain/replicate ≥ 1,544; * = p < 0.05, **** = p < 0.0001. (D) Expression of *K. lactis* Vrl1 (klVrl1-Envy) from the *RPL18B* promoter enhances klVps5-mScI localization in wild-type cells and induces Vps17-independent localization. (E) Quantification of RFP puncta per cell in *D*. Two tailed equal variance *t* tests; *n* = 4, cells/strain/replicate ≥ 144; ** = p < 0.01. Scale bars, 2 µm. mScI, mScarletI. OE, over-expressed. mScI, mScarletI. WGD, whole genome duplication.

To investigate if *K. lactis* Vps5 can form both VINE and retromer, we first generated an RFP-tagged *K. lactis* Vps5 construct for ectopic expression in *S. cerevisiae* (klVps5-mScI). We observed klVps5-mScI at small puncta in wild-type cells that were greatly enhanced in a *vps5*Δ strain and lost in a *vps17*Δ strain (p<0.0001 and p<0.05, respectively; Figure 4B, C), suggesting that klVps5 competes with *S. cerevisiae* Vps5 (scVps5) for scVps17 binding and retromer formation. Indeed, klVps5-mScI strongly co-purified with scVps29-3HA (Figure S3A) and partially rescued CPY sorting defects (Figure S3B). These results demonstrate that klVps5 can cooperate with the *S. cerevisiae* machinery to form a hybrid retromer complex with limited cargo sorting capabilities.

We then examined the ability of klVps5 to form a hybrid VINE complex. Neither scVrl1-Envy nor the artificially recruited scVrl1 chimera could recover the loss of klVps5-mScI puncta in *vin1*Δ*vps17*Δ cells (Figure S4A), but moderate recruitment of the klVps5 N-terminus by the artificially recruited scVrl1 chimera suggested that a weak interaction may occur despite species-specific differences (Figure S4B). We wondered if the *K. lactis* Vrl1 homolog (klVrl1) might more readily bind klVps5 to form VINE. Indeed, ectopic expression of klVrl1-Envy greatly increased bright klVps5-mScI puncta in wild-type cells and overcame the loss of puncta in *vin1*Δ*vps17*Δ cells (p<0.01; Figure 4D, E), suggesting that klVrl1 and klVps5 can cooperate to form the *K. lactis* version of VINE in *S. cerevisiae*, herein referred to as klVINE. Taken together our results indicate that klVps5 can form both klVINE and retromer.

### klVps5 recognizes klVrl1 and scVps29 at an overlapping site in its N-terminus

To understand how both VINE and retromer could assemble while sharing the single Vps5 isoform in *K. lactis*, we took a structural modeling approach. Using AlphaFold2, we first obtained confidently predicted interactions for the BAR domain of klVps5 with those of either klVrl1 or klVps17, consistent with the idea that klVps5 can participate in both complexes (Mirdita et al., 2022; Figure S5A, B). We then identified a region of the klVps5 N-terminus between aa 172-203 that was confidently predicted to associate with klVps29 (Figure 5A, Figure S6A). Like the interaction between *S. cerevisiae* Vps5 and Vps29 (Figure 3A), klVps5 was predicted to interact with klVps29 through bipartite pocket- and sheet-binding sites (Figure 5A). The pocket-binding sites of both klVps5 and scVps5 contain a common DPL motif, while the sheet-binding sites share the Leu-Phe motif which is predicted to interface with the same region of Vps29.

**Figure 5.**
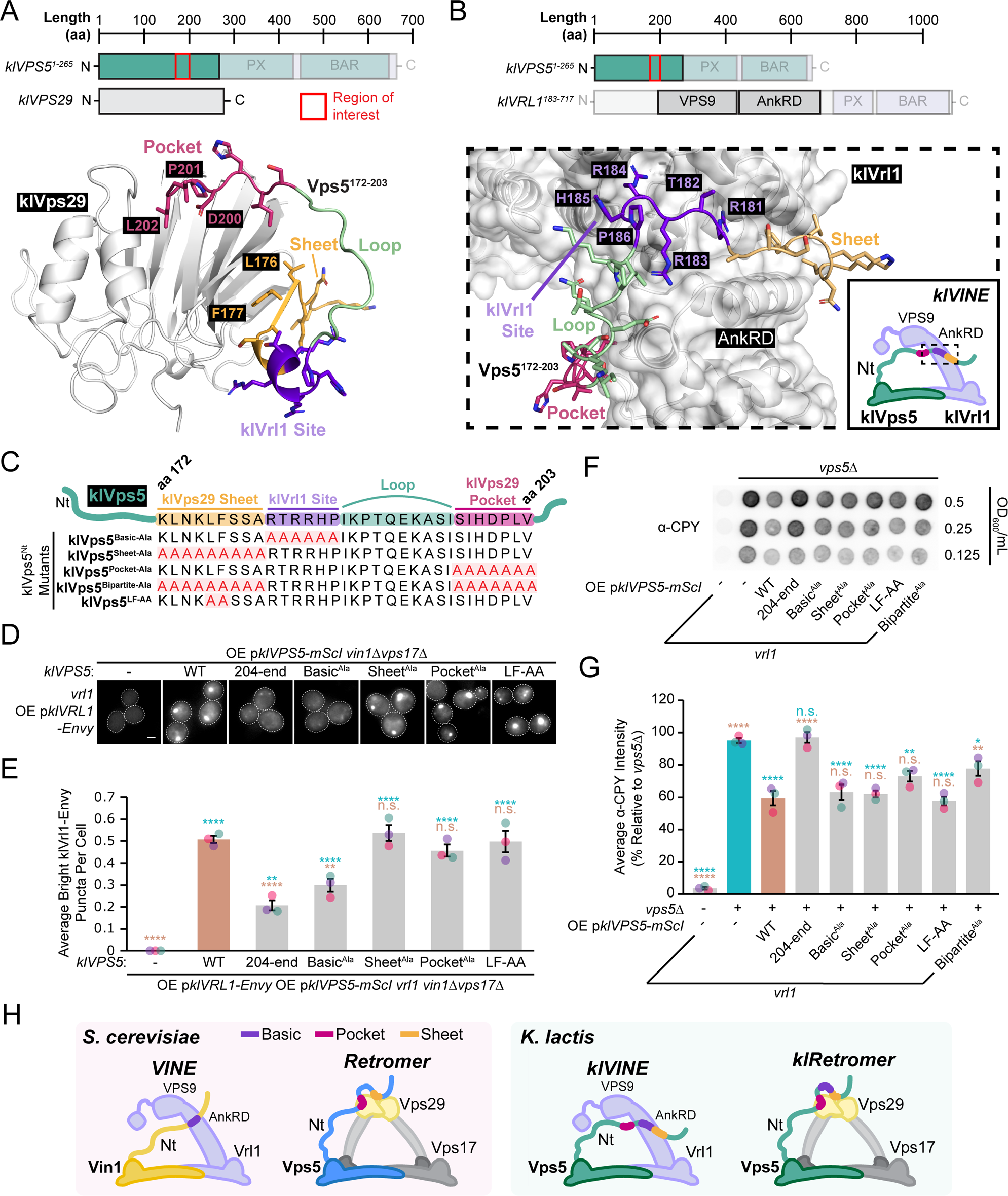
Nested motifs in the klVps5 N-terminus guide selection for retromer and VINE. (A) AlphaFold2-predicted interaction between klVps29 (white) and the klVps5 N-terminus (klVps5^1-204^). klVps5 is predicted to interact with klVps29 at two distinct sites, mirroring the analogous interaction in *S. cerevisiae*. klVps5 binds klVps29 through a bipartite motif between N-terminal residues aa 172-203 and involves a hydrophobic pocket site and a β-sheet site. (B) AlphaFold2-predicted interaction between klVrl1^183-717^ (white) and the klVps5 N-terminus (klVps5^1-204^). klVps5 interacts with klVrl1 in a similar manner to Vin1 and Vrl1 in *S. cerevisiae*, through a basic region involving the residues RTRRHP. (C) Diagram of klVps5 substitution mutations in the predicted N-terminal klVrl1-binding and klVps29 pocket- and sheet-binding sites generated to test VINE and retromer assembly. (D) *RPL18B*pr-driven expression of klVps5-mScI N-terminal mutants has differential effects on klVrl1-Envy localization in *vin1*Δ*vps17*Δ cells. Whereas deletion of the N-terminus (klVps5^204-end^) and alanine substitution of the polybasic predicted klVrl1-binding site causes loss of klVrl1 localization, disruption of either predicted klVps29 pocket- or sheet-binding sites have no effect. (E) Quantification of GFP puncta per cell in *D*. One-way ANOVA with Dunnett’s multiple comparison tests; *n* = 3, cells/strain/replicate ≥ 515; not significant, n.s. = p > 0.05, ** = p < 0.01, **** = p < 0.0001. Blue statistical significance labels correspond to a Dunnett-corrected ANOVA performed against the empty vector *vin1*Δ*vps17*Δ strain while brown labels correspond to a Dunnett-corrected ANOVA performed against wild-type klVps5-mScI. (F) Disruption of the klVps5 N-terminus either by complete deletion or alanine substitution of the pocket- and sheet-binding sites causes CPY secretion. Alanine substitution of the predicted klVrl1-binding polybasic site in the klVps5 N-terminus does not create a significant CPY secretion phenotype. (G) Quantification of secreted CPY by densitometry and normalized to measured secreted CPY from the *vps5*Δ strain. One-way ANOVA with Dunnett’s multiple comparison tests; n=3; not significant, n.s. = p > 0.05, * = p < 0.05, ** = p < 0.01, **** = p < 0.0001. Blue statistical significance labels correspond to a Dunnett-corrected ANOVA performed against a *vps5*Δ strain while brown labels correspond to a Dunnett-corrected ANOVA performed against a *vps5*Δ strain with plasmid-expressed wild-type klVps5-mScI. (H) Model for VINE and retromer assembly with N-terminal binding sites highlighted for Vin1 and Vps5, respectively. A polybasic region unique to Vin1 drives interaction with the Vrl1 AnkRD, while a bipartite motif in the Vps5 N-terminus associates at two distinct sites in Vps29—a conserved hydrophobic pocket and a β-sheet on the opposite surface of the protein. The sole Vps5 isoform in *K. lactis* possesses all three of these N-terminal motifs and can form both VINE and retromer. Error bars report SEM. aa, amino acid. mScI, mScarletI. Nt, N-terminus. OE, over-expressed.

Next, we used AlphaFold2 to predict binding between the entire klVps5 N-terminus and the VPS9 and AnkRD domains of klVrl1 (klVrl1^183-717^; Figure S6B). Strikingly, the ∼16 aa loop region between the klVps5 pocket- and sheet-binding sites contains a RTRRHP motif that was confidently predicted to interact with the same region of klVrl1 that forms the VINE interface in *S. cerevisiae* (Figure 5B, Figure S6B). This RTRRHP motif is not predicted to form any contacts with klVps29 and has electrochemical similarity to the polybasic region that we identified in *S. cerevisiae* Vin1 (Figure 1E, F). Taken together, these results suggest that the same ∼32 aa region of the klVps5 N-terminus contains overlapping signals that contribute to the formation of both VINE and retromer in *K. lactis*.

To test if these predicted interactions are important *in vivo*, we introduced a series of mutations to the N-terminus of klVps5-mScI (Figure 5C). klVrl1-Envy was largely displaced from bright puncta in *vin1*Δ*vps17*Δ cells when either the N-terminus of klVps5-mScI was truncated (klVps5^204-end^) or the basic RTRRHP motif was substituted to alanine (p<0.0001 and p<0.001, respectively; Figure 5D, E). In contrast, alanine substitution of either the predicted klVps29 sheet- and pocket-binding sites, or associated Leu-Phe motif, had no effect on VINE formation (Figure 5D, E), suggesting that the N-terminal polybasic site specifically contributes to ectopic VINE formation.

Despite only partial rescue of *S. cerevisiae vps5*Δ phenotypes by klVps5, mutations in the klVps5 N-terminus resulted in significant sorting defects. Truncation of the klVps5 N-terminus increased CPY secretion (Figure 5F, G), similar to the effect of deleting the *S. cerevisiae* Vps5 N-terminus (Figure 3G, H). Disruption of the pocket- and sheet-binding sites also caused a significant defect (p<0.01, Figure 5F, G). The effects of mutations in *K. lactis* Vps5 were in agreement with those in *S. cerevisiae* Vps5, with the bipartite mutations eliciting a more severe defect than mutations in either pocket- or sheet-binding sites alone, while not accounting for the full contribution of the N-terminal unstructured region (Figure 3G, H, Figure 5F, G). In contrast to the requirements for klVINE formation (Figure 5D, E), alanine substitution of the klVps5 N-terminal polybasic site did not result in obvious CPY sorting defects. Taken together, these data suggest that the N-terminus of *K. lactis* Vps5 may contribute to the formation of both VINE and retromer through interactions at a nested site (Figure 5H).

## Discussion

Here we revealed mechanisms underlying SNX-BAR assembly by studying the *S. cerevisiae* paralogs Vps5 and Vin1. We found that these proteins use shared and unique motifs in their N-termini to guide coat complex selection, with input from their respective BAR domains. Our results also suggest that nested motifs confer dual roles for the sole *K. lactis* Vps5 isoform as a subunit of both VINE and retromer. Taken together, our work delineates a mechanism by which endosomal coats select SNX-BAR proteins by recognizing unstructured N-terminal domains, which are common in this family of membrane adaptors.

### The N-termini of Vps5-family SNX-BARs promote coat assembly

The unstructured N-terminal regions of both Vin1 and Vps5 contribute to the specificity of endosomal coat formation in *S. cerevisiae*. Both Vin1 and Vps5 leverage a Leu-Phe motif, yet the Vin1 Leu-Phe motif is predicted to contact a Vrl1 hydrophobic patch, whereas the Vps5 Leu-Phe forms part of a short β-strand predicted to laterally extend a Vps29 β-sheet. Vin1 also interacts at a conserved surface on the Vrl1 AnkRD through a polybasic sequence, while a Vps5 FTDPL motif associates with Vps29 at a conserved hydrophobic pocket. Interestingly, the duplicated human orthologs of Vin1 and Vps5, SNX1/SNX2, also feature extended N-termini that interact with the retromer-associated sorting nexin SNX27 (Chandra et al. 2021; Simonetti et al., 2019; Yong et al., 2021). Yeast two-hybrid experiments have demonstrated that SNX1/SNX2 also bind to VPS29 and VPS35 (Haft et al., 2000; Rojas et al., 2007) and Gullapalli et al. (2004) mapped retromer binding to the SNX1 unstructured N-terminus.

Similarly to Vps5, SNX1 binds to the conserved hydrophobic pocket in VPS29 (Swarbrick et al., 2011). This pocket is of special interest since it is targeted by various effector proteins in humans (Baños-Mateos et al., 2019). The bacterial pathogen *L. pneumophila* interferes with retromer by binding the VPS29 pocket through its effector protein RidL (Finsel et al., 2013). Additionally, the Vrl1 homolog and RAB21 GEF VARP, the RAB7 GAP TBC1D5, the Commander complex subunit VPS35L, and the WASH complex subunit FAM21 also compete for this site (Crawley-Snowdon et al., 2020; Hesketh et al., 2014; Healy et al. 2023; Guo et al. 2023), suggesting that the VPS29 pocket is a multi-functional interface that could spatiotemporally control Rab dynamics and regulate interactions with sorting nexins and other accessory proteins. Further work is needed to determine the relevance of the interaction between SNX1 and the VPS29 pocket *in vivo*, and its role in regulating the incorporation of metazoan retromer into a variety of sorting complexes.

The unstructured N-termini of Vps5/Vin1 and SNX1/SNX2 are each at least one hundred aa long yet known interactions can be mapped to very short motifs (Simonetti et al., 2022). It is possible that remaining N-terminal sequences mediate additional regulatory interactions. SNX5/6 interact with p150^Glued^, a component of the dynactin machinery responsible for controlling dynein motor function, to drive cargo transport (Hong et al., 2009; Wassmer et al., 2009) and SNX2 binds the ER protein VAP to regulate lipid exchange at ER-endosome contacts (Dong et al., 2016). Accessory interactions could occur simultaneously to capture auxiliary factors that enhance sorting, or in a stepwise manner to ensure specificity while adding a layer of temporal control.

### Assembly and function of VINE and retromer in *K. lactis*

The genome of milk yeast *K. lactis* encodes individual copies of the *VPS5*, *VPS17* and *VRL1* genes. Our experiments based on ectopic expression in *S. cerevisiae* suggest that klVps5 forms both VINE and retromer, at least in part through nested motifs in its N-terminus. The use of overlapping signals ensures that Vps5 can engage with only one complex at a time, and raises questions about how the relative levels of retromer and VINE are regulated. The partitioning of klVps5 between retromer and VINE complexes could be driven by its abundance and its relative affinity for klVrl1 or subunits of retromer. In theory, post-translational modifications could alter the activity or availability of one motif. Because the relevant pathways are unlikely to be present in *S. cerevisiae*, further studies in *K. lactis* are needed to uncover the regulatory mechanisms that govern subunit sharing. Taken together, our work has clarified mechanisms of SNX-BAR complex assembly in yeast, which could provide insight into similar strategies used by human SNX-BARs.

## Materials and Methods

| Key Resources Table |  |  |  |  |
| --- | --- | --- | --- | --- |
| Reagent type (species) or resource | Designation | Source or reference | Identifiers | Additional information |
| Antibody | Anti-HA (Mouse monoclonal) | Covance | MMS-101R; HA.11 | WB (1:1,000) |
| Antibody | Anti-RFP (Mouse monoclonal) | ChromoTek | 6G6 | WB (1:2,000) |
| Antibody | Anti-HA (Rabbit polyclonal) | AbCam | Ab9110 | CoIP |
| Antibody | HRP-Anti-Mouse (Goat polyclonal) | Jackson | 115-035-146 | WB (1:20,000) |
| Antibody | Anti-CPY (Mouse monoclonal) | Thermo Fisher Scientific | A-6428 | WB (1:500) |
| Other | nProtein A Sepharose 4 Fast Flow | Cytiva | 17528004 | CoIP |
| Other | Amersham ECL | Cytiva | GERPN2209 | Chemiluminescent reagent |
| Other | Amersham ECL Prime | Cytiva | GERPN2232 | Chemiluminescent reagent |
| Other | Yeast/Fungal ProteaseArrest | GBiosciences | 786-435 | Yeast protease inhibitor; 100X |
| Other | Concanavalin A | Sigma-Aldrich | C2010 | Yeast live-cell imaging preparation |
| Software, algorithm | MetaMorph | MDS Analytical Technologies | Version 7.8 | Automated image analysis |
| Software, algorithm | GraphPad Prism | GraphPad Software | Version 9.1.0 | Statistical analysis |
| Software, algorithm | ColabFold<br>AlphaFold2<br>Advanced | Jumper et al., 2021; Mirdita et al., 2022 |  | Protein structure and binding prediction software |

### Yeast strains and plasmids

Yeast strains and plasmids used in this study are described in Supplementary Files 1 and 2, respectively. Yeast strains were built in the BY4741 strain background using homologous recombination-based integration unless otherwise indicated. Gene deletions, promoter exchanges and tags were confirmed by colony PCR and either western blot or fluorescence microscopy where possible. Plasmids were built by NEBuilder® HiFi DNA assembly or homologous recombination in yeast, recovered in *Escherichia coli*, and confirmed by sequencing. *K. lactis* DNA constructs were built using native sequences obtained from strain NRRL Y-1140 (ATCC 8585).

### Bioinformatic analysis of protein folding

Prediction of protein structure and binding interfaces was performed using the AlphaFold2 advanced server accessed through the ColabFold platform with default settings (Jumper et al., 2021; Mirdita et al., 2022). Structural models were processed for presentation using PyMOL2 (Schrödinger, LLC, New York, New York).

### Fluorescence microscopy and automated image analysis

Yeast strains were prepared for microscopy by overnight growth, followed by four hours of growth at 30 °C in fresh synthetic dextrose-based (SD) media and plating on 96-well glass-bottom plates (Cellvis, Mountain View, California) coated in concanavalin A. Images were collected using a DMi8 microscope (Leica Microsystems, Wetzlar, Germany) equipped with an ORCA-flash 4.0 digital camera (Hamamatsu Photonics, Shizuoka, Japan) and a high-contrast Plan-Apochromat 63x/1.30 Glyc CORR CS oil immersion lens (Leica Microsystems, Wetzlar, Germany). The MetaMorph 7.8 software package was used for image acquisition and processing (MDS Analytical Technologies, Sunnyvale, California). The intensity of a given fluorophore is scaled identically in all micrographs within an experiment. Images were resized using Photoshop CC 2020 (Adobe, San Jose, California) and arranged in Illustrator CC 2020 (Adobe, San Jose, California). MetaMorph 7.8 scripted journals were used to quantify raw images (MDS Analytical Technologies, Sunnyvale, California). The Count Nuclei feature utilizing intensity above local background (IALB) was used to exclude dead cells and identify live cells. The Granularity feature was used to identify puncta in a dead cell-masked intermediate image based on IALB. Masking functions were performed using the Arithmetic function with Logical AND. RFP channel signal is represented as magenta in merged micrographs to improve accessibility.

### Coimmunoprecipitation and western blotting

For co-immunoprecipitation analyses, yeast cultures were grown to log phase at 30 °C in synthetic dextrose-based (SD) media and 75 OD_600_ of yeast were collected and incubated for 15 minutes in 50 mM Tris-Cl with 10 mM DTT (pH 9.5) at room temperature, followed by one hour digestion in spheroplasting buffer (1.2 M sorbitol, 50 mM KH2PO4, 1 mM MgCl2 and 250 µg/ml zymolase at pH 7.4) at 30 °C. After washing spheroplasts twice with 1.2 M sorbitol, they were frozen at −80°C then incubated for 10 minutes at room temperature in 500 µl lysis buffer (0.5% Triton X-100, 50 mM HEPES, 1 mM EDTA, 50 mM NaCl, 1 mM PMSF and 1x fungal Protease Arrest, pH 7.4). 50 µl of lysate was collected for each sample and mixed with 2x Laemmli buffer (4% SDS, 20% glycerol, 120 mM Tris-Cl (pH 6.8), 0.01g bromophenol blue and 10% beta-mercaptoethanol) for analysis by western blot. A polyclonal rabbit anti-HA antibody (ab9110, Abcam) was added to remaining lysates and incubated at 4 °C for 1 hour. Protein A Sepharose beads (Cytiva) were added and incubated at 4 °C for 1 hour, then beads were washed three times in lysis buffer and resuspended in 50 µl of Thorner buffer (8 M Urea, 5% SDS, 40mM Tris-Cl (pH 6.4), 1% beta-mercaptoethanol, 0.4 mg/mL bromophenol blue) and heated for five minutes at 80 °C. Proteins were separated on 8% SDS-PAGE gels and detected by western blot using monoclonal mouse anti-HA (MMS-101R; Covance) or monoclonal mouse anti-RFP antibodies (6G6; ChromoTek) prior to secondary antibody treatment with polyclonal goat anti-mouse conjugated to horseradish peroxidase (115–035-146; Jackson ImmunoResearch Laboratories). Blots were developed with Amersham ECL (GERPN2209, Cytiva) or Amersham ECL Prime (GERPN2232, Cytiva) chemiluminescent western blot detection reagents and exposed using the Vilber Fusion FX with automatic settings (Vilber Smart Imaging, Collégien, France). Densitometry of scanned films was performed using ImageJ (Schneider et al., 2012).

For detection of secreted carboxypeptidase Y (CPY), yeast cultures were grown to log phase at 30 °C, serially diluted 1:1 and spotted to synthetic dextrose (SD) based media and incubated 12-15 hours at 30 °C under a nitrocellulose membrane. For test of CPY secretion in klVps5-expressing strains the *vps5*Δ control strain was spotted in duplicate to provide separate data points for normalization and statistical comparison. Nitrocellulose was washed with dH2O and probed using a monoclonal mouse anti-CPY antibody (A-6428, Thermo Fisher Scientific) followed by secondary antibody treatment with polyclonal goat anti-mouse conjugated to horseradish peroxidase (115–035-146; Jackson ImmunoResearch Laboratories) and development as described above.

### Statistical analysis of quantitative data

Statistical analyses were performed using GraphPad Prism 9.1.0 (GraphPad Software, San Diego, California). A 95% confidence threshold (P < 0.05) was used for hypothesis testing. Microsoft Excel 2019 was used to graph data (Microsoft, Redmond, Washington). Bar graphs show mean value of all collected biological replicates, with data points from individual replicates represented as scatter plots coloured by replicate. Error bars represent standard error of the mean (SEM).

## Acknowledgements

We thank Dr. Luc Berthiaume (University of Alberta, Edmonton, Canada) for generously sharing rabbit anti-GFP serum. We gratefully acknowledge funding support from the Natural Sciences and Engineering Research Council of Canada (grant RGPIN-2022-04573 to EC and PGS-D3 Doctoral Scholarship to SPS); Canada Foundation for Innovation (Leading Edge Fund 30636); Canadian Institutes of Health Research (CGS-M Frederick Banting and Charles Best Canada Graduate Scholarship to SPS and MSF and CGS-D Frederick Banting and Charles Best Canada Graduate Scholarship to MSF); BC Children’s Hospital Research Institute Jan M. Friedman Graduate Studentship to SPS; University of British Columbia Catalyst Paper Corporation Affiliated Fellowship to SPS, Gertrude Langridge Graduate Scholarship and Elwyn Gregg Memorial Affiliated Fellowships to MSF; 4-Year Doctoral Fellowship to SPS and MSF; and University of British Columbia Medical Genetics Rotation Award to MSF.

## Supplementary Figure and File Legends

**Figure S1. Confidence measures of Vps29-Vps5 N-terminus binding prediction.** (A) AlphaFold2-predicted interaction between Vps29 and the N-terminus of Vps5 (Vps5^1-276^) with pLDDT scores mapped to each residue of Vps29 (top) or Vps5 (bottom). In each case, the interacting partner is shown in gray. (B) AlphaFold2-generated predicted alignment error (PAE; Jumper et al., 2021; Mirdita et al., 2022) for Vps29 and Vps5^1-276^ indicates an intermolecular interaction in the off-diagonal boxes. aa, amino acids.

**Figure S2. The SNX-BAR domains of Vin1 and Vps5 are indispensable for endosomal coat targeting.** (A) Schematic of Vin1 and Vps5 truncations and chimeric fusion constructs used in *B, C*. (B) The full-length Vin1-mScI protein is required to complement the loss of Vrl1-Envy puncta in *vin1*Δ cells. A chimeric protein with the N-terminus of Vin1 and SNX-BAR domains of Vps5 also fails to complement in both *vin1*Δ and *vin1*Δ*vps17*Δ cells. (C) The SNX-BAR domains of Vps5 are sufficient to partially complement the loss of Vps35-GFP puncta in *vps5*Δ cells. A chimeric protein with the N-terminus of Vps5 and SNX-BAR domains of Vin1 fails to complement in both *vps5*Δ and *vps5*Δ*vps17*Δ cells.

**Figure S3. klVps5 binds Vps29 and partially complements CPY secretion in *S. cerevisiae*.** (A) Both mScI-tagged *S. cerevisiae* and *K. lactis* isoforms of Vps5 co-purify with Vps29-3HA in *S. cerevisiae*. (B) Both mScI-tagged *S. cerevisiae* and *K. lactis* isoforms of Vps5 partially complement the CPY secretion in *vps5*Δ *S. cerevisiae* cells. For the *S. cerevisiae* Vps5 isoform, complementation depends on its unstructured N-terminus.

**Figure S4. klVps5 does not readily form VINE with scVrl1.** (A) *S. cerevisiae* Vrl1-Envy and *K. lactis* Vps5-mScI both fail to localize in *vin1*Δ*vps17*Δ cells. The artificially recruited Vrl1(1-703)^YPE^ chimera does not recruit *K. lactis* Vps5-mScI to puncta. (B) The artificially recruited Vrl1(1-703)^YPE^ chimera weakly recruits the *K. lactis* Vps5 N-terminus fused to mScI in *vin1*Δ cells. mScI, mScarletI. YPE, Ypt35(PX)-Envy.

**Figure S5. klVps5 forms predicted BAR dimers with klVps17 and klVrl1.** (A) AlphaFold2-predicted interaction between the *K. lactis* Vps5 and Vps17 BAR domains along the canonical BAR-BAR dimerization interface (top) and with pLDDT scores mapped to each residue (bottom). (B) AlphaFold2-predicted interaction between the *K. lactis* Vps5 and Vrl1 BAR domains along the canonical BAR-BAR dimerization interface (top) and with pLDDT scores mapped to each residue (bottom).

**Figure S6. Confidence measures of klVps5 N-terminal binding predictions with klVrl1 or klVps29.** (A) AlphaFold2-predicted interaction between *K. lactis* Vps29 and the N-terminus of *K. lactis* Vps5 (klVps5^1-265^) with pLDDT scores mapped to each residue of Vps29 (bottom left) or Vps5 (bottom right). In each case, the interacting partner is shown in gray. AlphaFold2-generated predicted alignment error (PAE; Jumper et al., 2021; Mirdita et al., 2022) for klVps29 and klVps5^1-265^ indicates an intermolecular interaction in the off-diagonal boxes (top right). (B) AlphaFold2-predicted interaction between the *K. lactis* Vrl1 VPS9 and AnkRD domains (klVrl1^183-717^) and the N-terminus of *K. lactis* Vps5 (klVps5^1-265^) with pLDDT scores mapped to each residue of klVrl1 (bottom left) or Vps5 (bottom right). In each case, the interacting partner is shown in gray. AlphaFold2-generated predicted alignment error (PAE; Jumper et al., 2021; Mirdita et al., 2022) for klVrl1^183-717^ and klVps5^1-265^ indicates an intermolecular interaction in the off-diagonal boxes (top right). aa, amino acids.

**Supplementary File 1. List of *Saccharomyces cerevisiae* strains used in this study.**

**Supplementary File 2. List of plasmids used in this study.**

