## Supplementary Figures S1-S6 and Supplementary Files 1 and 2 for "N-terminal signals in the SNX-BAR paralogs Vps5 and Vin1 guide coat complex formation"

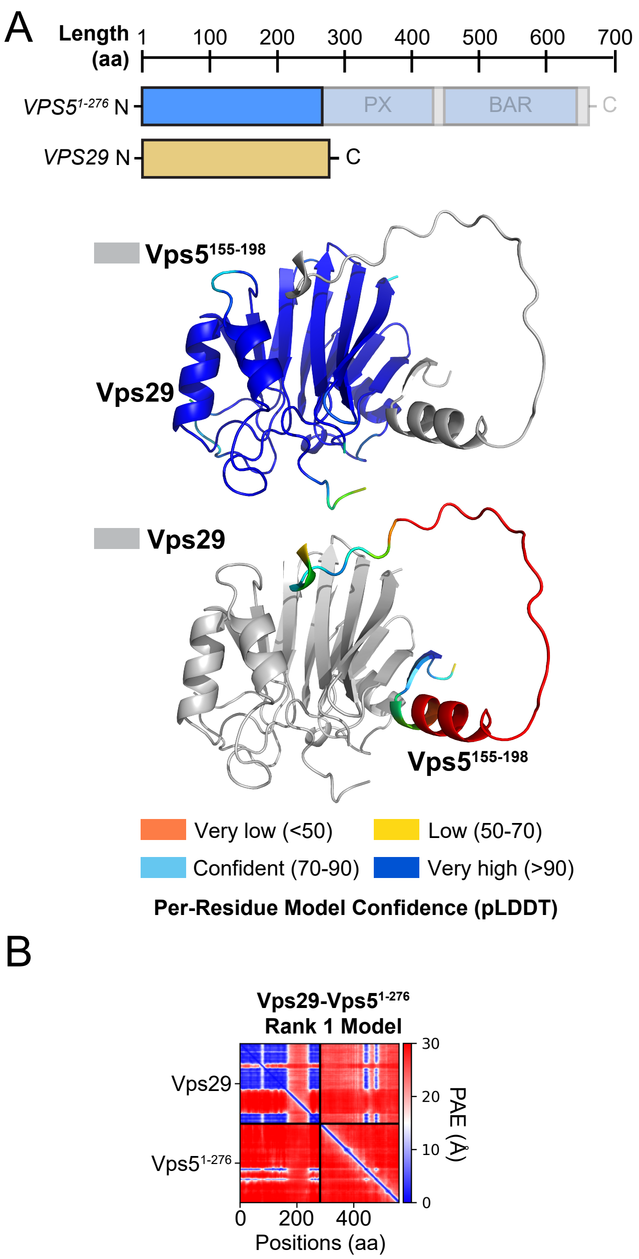


**Figure S1,** Shortill et al., 2024

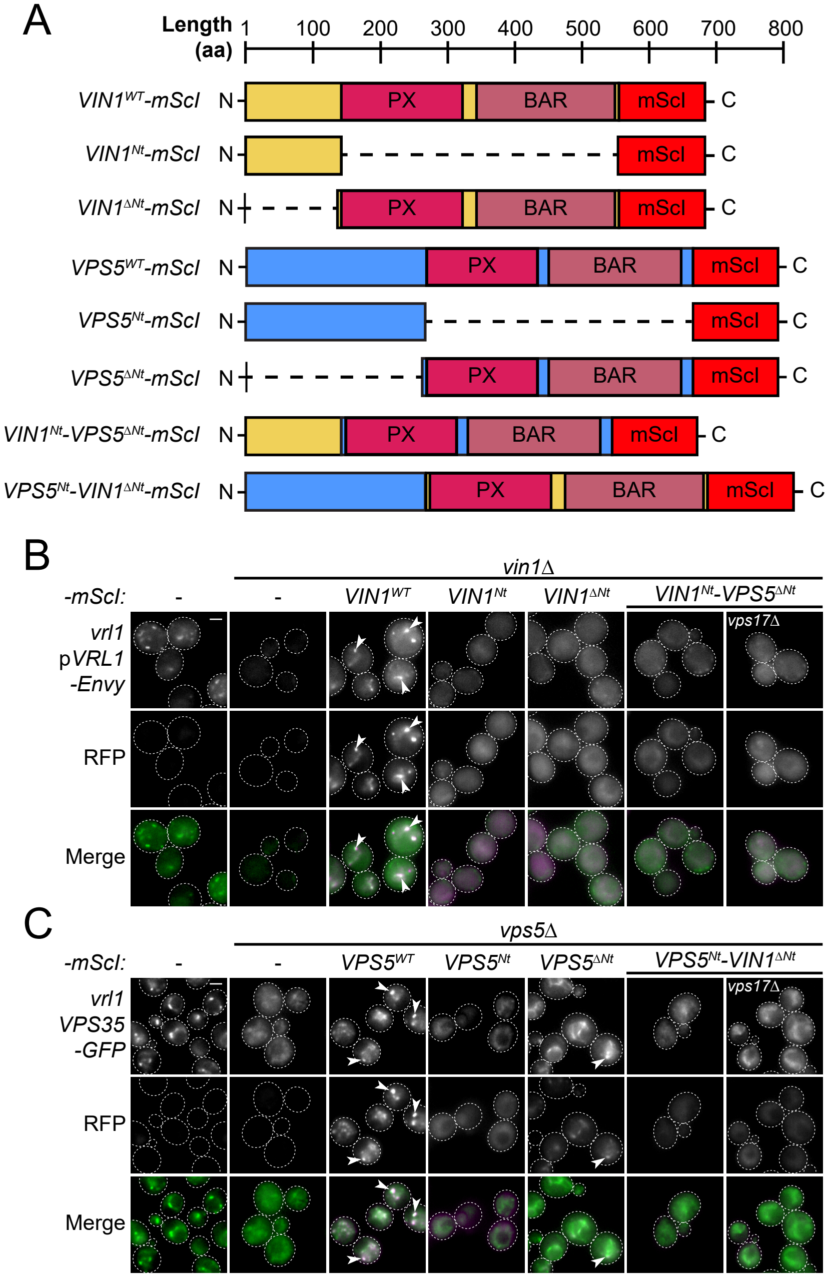


**Figure S2,** Shortill et al., 2024

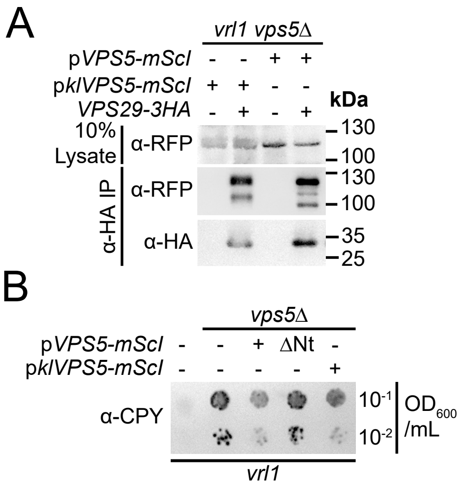


**Figure S3,** Shortill et al., 2024

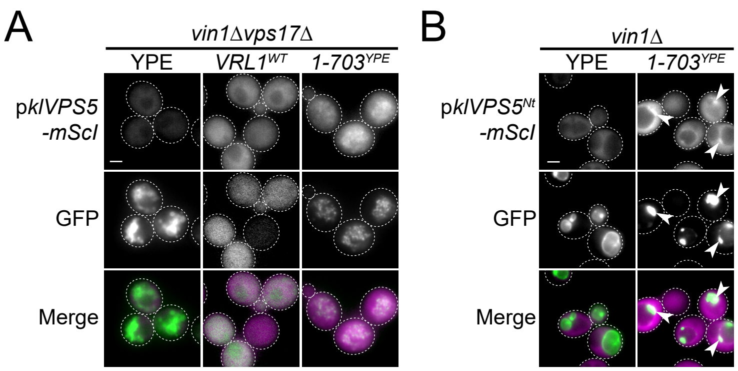


**Figure S4,** Shortill et al., 2024

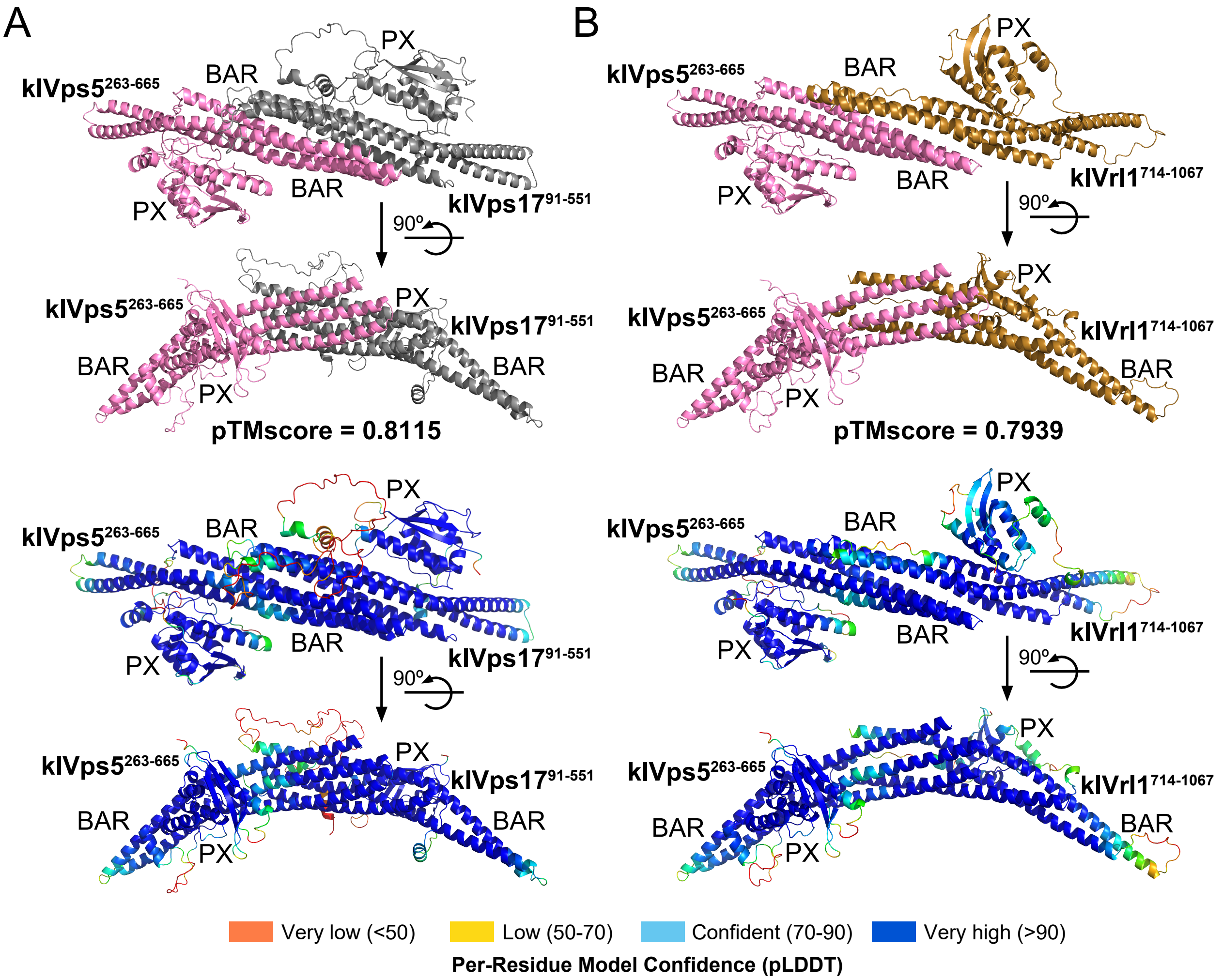


**Figure S5,** Shortill et al., 2024

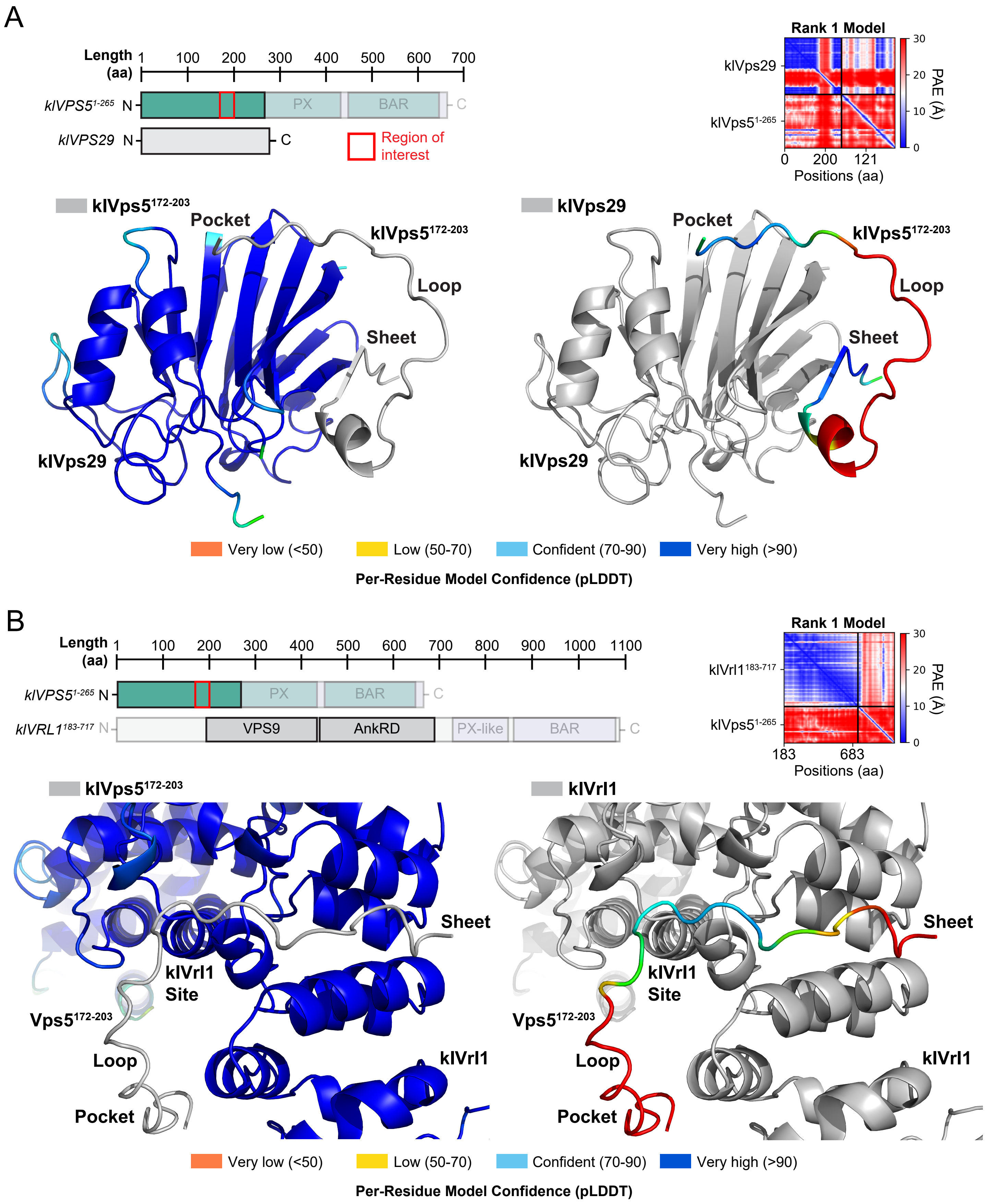


**Figure S6,** Shortill et al., 2024

| **Table S1: Yeast strains used in this study** | | |
|---|---|---|
| **Strain Name** | **Genotype** | **Source** |
| BY4741 | *MATa his3Δ1 leu2Δ0 met15Δ0 ura3Δ0* | Dr. EK O'Shea (Harvard), Huh et al., 2003 (PMID: 14562095) |
| *vin1Δ* | BY4741 *vin1Δ::KAN* | MATa Deletion Mutant Collection, Giaever et al., 2002 (PMID: 12140549) |
| SSY096 | BY4741 *vps5Δ::NAT* | Shortill et al., 2022 (PMID: 35938928) |
| SSY1063 | BY4741 *VPS29-3HA::KAN vps5Δ::NAT* | This Study |
| SSY1029 | BY4741 *vin1Δ::KAN vps17Δ::Hph* | This Study |
| *VPS35-GFP* | BY4741 *VPS35-GFP::HIS3* | Dr. EK O'Shea (Harvard), Huh et al., 2003 (PMID: 14562095) |
| SSY097 | BY4741 *VPS35-GFP::HIS3 vps5Δ::NAT* | Shortill et al., 2022 (PMID: 35938928) |
| SSY730 | BY4741 *VPS35-GFP::HIS3 vps17Δ::Hph* | Shortill et al., 2022 (PMID: 35938928) |
| SSY732 | BY4741 *VPS35-GFP::HIS3 vps5Δ::NAT vps17Δ::Hph* | Shortill et al., 2022 (PMID: 35938928) |

| Table S2: Plasmids used in this study | | | |
|---|---|---|---|
| **Plasmid Name** | **Genotype** | **Marker** | **Source** |
| pRS415 | pRS415 CEN *AmpR LEU2* | *LEU2* | Sikorski and Hieter, 1989 (PMID: 2659436) |
| pRS416 | pRS416 CEN *AmpR URA3* | *URA3* | Sikorski and Hieter, 1989 (PMID: 2659436) |
| pMF24 | pRS416-*ADH1pr-YPT35(61-end)-Envy* | *URA3* | This Study |
| pSS052 | pRS416-*ADH1pr-VRL1(1-703)-YPT35(61-end)-Envy* | *URA3* | Shortill et al., 2022 (PMID: 35938928) |
| pMD454 | pRS415-*VIN1pr-VIN1-mScarletI* | *LEU2* | Shortill et al., 2022 (PMID: 35938928) |
| pMD533 | pRS415-*VIN1pr-VIN1(76-77Ala)-mScarletI* | *LEU2* | This Study |
| pMD534 | pRS415-*VIN1pr-VIN1(78-79Ala)-mScarletI* | *LEU2* | This Study |
| pMD535 | pRS415-*VIN1pr-VIN1(80-81Ala)-mScarletI* | *LEU2* | This Study |
| pMD536 | pRS415-*VIN1pr-VIN1(82-83Ala)-mScarletI* | *LEU2* | This Study |
| pMD537 | pRS415-*VIN1pr-VIN1(84-85Ala)-mScarletI* | *LEU2* | This Study |
| pMD538 | pRS415-*VIN1pr-VIN1(86-87Ala)-mScarletI* | *LEU2* | This Study |
| pMD539 | pRS415-*VIN1pr-VIN1(88-89Ala)-mScarletI* | *LEU2* | This Study |
| pMD540 | pRS415-*VIN1pr-VIN1(90-91Ala)-mScarletI* | *LEU2* | This Study |
| pMD541 | pRS415-*VIN1pr-VIN1(92-93Ala)-mScarletI* | *LEU2* | This Study |
| pMD542 | pRS415-*VIN1pr-VIN1(94-95Ala)-mScarletI* | *LEU2* | This Study |
| pSS100 | pRS415-*VIN1pr-VIN1(83-84Ala)-mScarletI* | *LEU2* | This Study |
| pSS059 | pRS416-*VIN1pr-VIN1(1-116)-mScarletI* | *URA3* | Shortill et al., 2022 (PMID: 35938928) |
| pSS055 | pRS415-*ADH1pr-YPT35(61-end)-Envy* | *LEU2* | Shortill et al., 2022 (PMID: 35938928) |
| pSS046 | pRS415-*ADH1pr-VRL1(1-703)-YPT35(61-end)-Envy* | *LEU2* | Shortill et al., 2022 (PMID: 35938928) |
| pSS077 | pRS415-*ADH1pr-VRL1(1-703-L497D)-YPT35(61-end)-Envy* | *LEU2* | This Study |
| pPW1 | pRS416-*VRL1pr-VRL1-Envy* | *URA3* | Shortill et al., 2022 (PMID: 35938928) |
| pMD552 | pRS416-*VPS5pr-VPS5-mScarletI* | *URA3* | This Study |
| pMD571 | pRS416-*VPS5pr-VPS5(275-end)-mScarletI* | *URA3* | This Study |
| pMD560 | pRS416-*VPS5pr-VPS5(Pocket-Ala)-mScarletI* | *URA3* | This Study |
| pMD556 | pRS416-*VPS5pr-VPS5(Sheet-Ala)-mScarletI* | *URA3* | This Study |
| pMD559 | pRS416-*VPS5pr-VPS5(Bipartite-Ala)-mScarletI* | *URA3* | This Study |
| pMD562 | pRS416-*VPS5pr-VPS5(LF-AA)-mScarletI* | *URA3* | This Study |
| pMD561 | pRS416-*VPS5pr-VPS5(L196K)-mScarletI* | *URA3* | This Study |
| pMF10 | pRS415-*VRL1pr-VRL1-Envy* | *LEU2* | Shortill et al., 2022 (PMID: 35938928) |
| pSS056 | pRS416-*VIN1pr-VIN1-mScarletI* | *URA3* | Shortill et al., 2022 (PMID: 35938928) |
| pSS080 | pRS416-*VIN1pr-VIN1(111-end)-mScarletI* | *URA3* | This Study |
| pSS075 | pRS416-*VIN1pr-VIN1(111-end)-VPS5(275-end)-mScarletI* | *URA3* | This Study |
| pSS071 | pRS416-*VIN1pr-VPS5-mScarletI* | *URA3* | This Study |
| pSS060 | pRS416-*VIN1pr-VPS5(1-276)-mScarletI* | *URA3* | Shortill et al., 2022 (PMID: 35938928) |
| pSS074 | pRS416-*VIN1pr-VPS5(275-end)-mScarletI* | *URA3* | This Study |
| pSS073 | pRS416-*VIN1pr-VPS5(1-276)-VIN1(111-end)-mScarletI* | *URA3* | This Study |
| pMD510 | pRS416-*VIN1pr-klVPS5-mScarletI* | *URA3* | This Study |
| pSS093 | pRS415-*VRL1pr-klVRL1-Envy* | *LEU2* | This Study |
| pMD554 | pRS416-*RPL18Bpr-klVPS5-mScarletI* | *URA3* | This Study |
| pSS103 | pRS415-*RPL18Bpr-klVRL1-Envy* | *LEU2* | This Study |
| pMD565 | pRS416-*RPL18Bpr-klVPS5(204-end)-mScarletI* | *URA3* | This Study |
| pMD573 | pRS416-*RPL18Bpr-klVPS5(Basic-Ala)-mScarletI* | *URA3* | This Study |
| pMD566 | pRS416-*RPL18Bpr-klVPS5(Sheet-Ala)-mScarletI* | *URA3* | This Study |
| pMD567 | pRS416-*RPL18Bpr-klVPS5(Pocket-Ala)-mScarletI* | *URA3* | This Study |
| pMD568 | pRS416-*RPL18Bpr-klVPS5(LF-AA)-mScarletI* | *URA3* | This Study |
| pMF66 | pRS416-*RPL18Bpr-klVPS5(Bipartite-Ala)-mScarletI* | *URA3* | This Study |
